# The core microbiome as a reproducible, data-driven abstraction, not a biological entity

**DOI:** 10.64898/2026.09.15.751703

**Authors:** Camilo Quiroga-González, Luis Daniel Prada-Salcedo, Kezia Goldmann

**Affiliations:** Department of Agroecosystem Ecology, UFZ - Helmholtz Centre for Environmental Research, Theodor-Lieser-Straße 4, 06120, Halle (Saale), Germany; Department of Applied Microbial Ecology, UFZ - Helmholtz Centre for Environmental Research, Permoserstraße 16, 04318 Leipzig, Germany

**Keywords:** core microbiome, ecological consistency, ITS2, prevalence, plant-associated microbiome, *Quercus robur*, 16S rRNA

## Abstract

Microbial communities are highly diverse, making it difficult to distinguish stable ecological patterns from stochastic variation. The core microbiome offers a widely used framework for simplification, but prevalence-based definitions are often criticized for relying on arbitrary thresholds. We propose an alternative view in which the core microbiome is not a biologically complete or functionally exhaustive subset, but a data-driven abstraction that preserves dominant ecological information under strong dimensional reduction. Using bacterial (16S rRNA) and fungal (ITS2) communities from leaves, roots, and rhizosphere of clonally replicated pedunculate oak (*Quercus robur*) across a continental environmental gradient, we defined cores using a non-arbitrary threshold derived from the prevalence distribution. Despite retaining fewer than 6% of bacterial and 3% of fungal OTUs, cores preserved composition patterns, reproduced site differentiation, retained most predicted functional information, maintained network structure, and were robust across a wide range of sampling efforts. Prevalence and abundance captured distinct ecological dimensions: consistently occurring taxa were not necessarily the most abundant, and vice versa. Our results show that core value therefore lies not in identifying the most important microorganisms, but in providing a reproducible, information-preserving representation of complex communities — though core taxa nonetheless retained fundamental ecological and functional roles.

## Introduction

Microbial communities are among the most complex biological systems studied in ecology, characterized by extraordinary richness, high turnover, and strong context dependency across space and environmental gradients (Hanson *et al*., 2012; Gibbons & Gilbert, 2015; Bäcker *et al*., 2026). Although high-throughput sequencing provides unprecedented resolution for characterizing these communities, it often results in increasingly sparse and high-dimensional datasets that are difficult to interpret and compare across systems (Knight *et al*., 2018). Consequently, reducing community complexity while preserving ecological meaningful information has become a central challenge in microbial ecology.

A widely used approach to address this challenge is the concept of the “core microbiome”, commonly defined as the set of microbial taxa consistently detected across a majority of samples within a given system (Turnbaugh et al., 2007; Hamady & Knight, 2009; Shade & Handelsman, 2012; Neu et al., 2021). By definition, the core microbiome is thus mainly prevalence-based. Because these taxa are repeatedly observed across individuals or environments, the core microbiome is often assumed to represent the most ecologically stable, persistent, or potentially most important members of a microbial community (Turnbaugh *et al*., 2007; Shade & Handelsman, 2012). However, despite its widespread use, both, the conceptual interpretation and analytical implementation of the core microbiome remains highly inconsistent among studies (Lemanceau *et al*., 2017; Risely, 2020).

One major source of this inconsistency is that prevalence-based core definitions typically rely on arbitrary prevalence thresholds, which vary widely between studies and can substantially alter which taxa are considered part of the core (Ainsworth *et al*., 2015; Risely *et al*., 2021; Lee *et al*., 2026). Importantly, this limitation concerns the arbitrary selection of prevalence thresholds rather than the use of prevalence itself. Prevalence provides a meaningful measure of ecological consistency, but only when thresholds are determined objectively rather than chosen *ad hoc*. Recent studies have attempted to address this by identifying core taxa through occupancy-abundance approaches, which rank taxa by how consistently and abundantly they occur across samples and by their contribution to Bray-Curtis dissimilarity between samples, rather than relying on a single prevalence cutoff (Shade & Stopnisek, 2019; Lee et al., 2026). However, these studies also show that different thresholds applied to the same dataset yield different results, indicating that even such more objective approaches still require a standardized, dimensionality-reduction-based strategy. Beyond threshold selection, prevalence-based definitions implicitly prioritizes taxa that are consistently detected across samples, thereby favoring ecological generalists while excluding specialists that may be locally abundant but spatially restricted (Huse *et al*., 2012; Jousset *et al*., 2017; Chakraborty *et al*., 2025).

These limitations have motivated the development of alternative conceptual frameworks that redefine the core microbiome according to ecological function or microbial interactions rather than prevalence alone. Such approaches identify groups of microorganisms by functional traits, dominance patterns, co-occurrence networks, or inferred contributions to ecosystem processes (Shade & Handelsman, 2012; Risely, 2020).

However, these different approaches may identify microbial groups that do not necessarily overlap with prevalence-based cores, reinforcing the idea that different criteria capture distinct and partially non-overlapping aspects of microbial community structure (Fig. 1a). Consequently, the core microbiome should not be viewed as a universally defined biological entity.

**Fig. 1.**
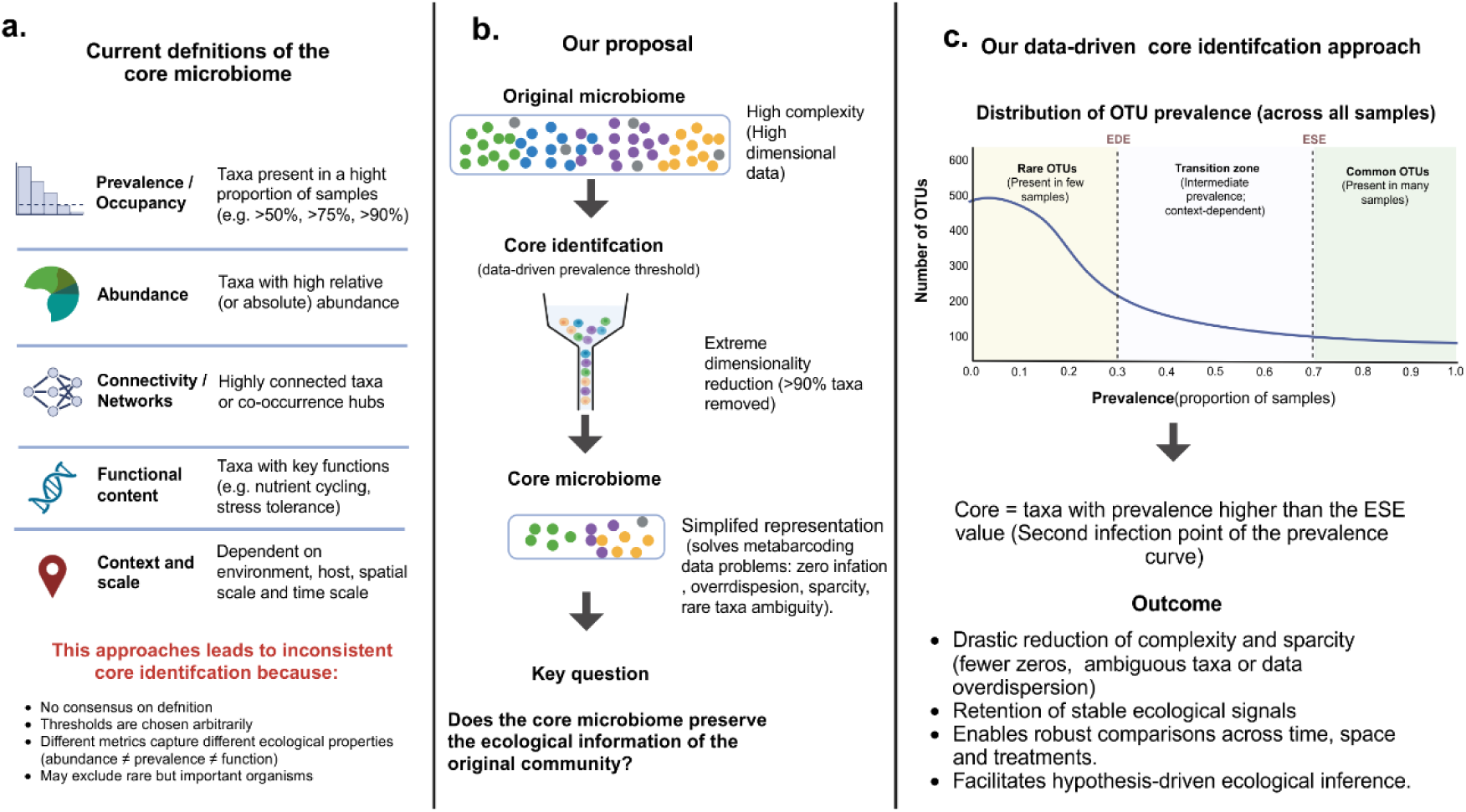
A data-driven framework for identifying the core microbiome. (a) Current criteria used to define core microbiomes, including prevalence, abundance, network position, functional relevance, and environmental context, which may lead to inconsistent definitions due to arbitrary thresholds and differences in ecological properties captured. (b) Conceptual overview of our approach: dimensional microbial community data are reduced into a core representation using a data-driven prevalence threshold, with the aim of retaining ecological information while reducing complexity and sparsity. (c) Identification of the core microbiome from the prevalence distribution of OTUs across samples. The core is defined as taxa exceeding the second inflection point of the curve (ESE). This threshold separates common, stable taxa from the transition zone and rare taxa, providing a reproducible framework for comparative ecological analyses (Created in BioRender).

This distinction raises a key conceptual issue: abundance, prevalence, functional importance, and ecological relevance are not equivalent properties of microbial communities (Lynch & Neufeld, 2015; Jousset *et al*., 2017; Banerjee *et al*., 2018). Highly abundant taxa may be spatially restricted, while low-abundance taxa may be widely distributed. Similarly, microbial taxa that are functionally important in specific contexts may be excluded from prevalence-based cores due to their limited occupancy. These inconsistencies suggest that the core microbiome should not be interpreted as a biologically complete or functionally exhaustive community subset. Thus, prevalence alone cannot be interpreted as a direct measure of ecological importance. Instead, prevalence primarily reflects ecological consistency. This suggests a different interpretation of the core microbiome. Rather than viewing the core as a biologically complete or functionally exhaustive subset of a microbial community, we propose that it is more appropriately understood as an analytical abstraction. Under this view, prevalence-based core identification represents a form of structured taxonomic filtering that reduces dataset dimensionality that aims to retain stable ecological signal while reducing stochastic variation and dataset complexity. The relevant question is therefore no longer which microorganisms belong to the core, but rather whether the reduced community preserves meaningful multivariate ecological structure in a simplified representation of the original community.

To evaluate this conceptual framework, we tested whether a prevalence-based core identified using an objective, data-driven prevalence threshold preserves ecological information despite massive reductions in taxonomic diversity (Fig. 1). We applied this approach to a large multi-compartment plant-associated microbiome dataset sampled from replicated clonal oak trees planted across 12 sites spanning a broad environmental gradient from France to Finland, encompassing land-use types ranging from grasslands to forests, with microbial communities sampled from the rhizosphere, roots, and leaves of each tree. Because microbial communities differ strongly among host-associated compartments, which represent distinct ecological and selective environments (Bais *et al*., 2006; Lundberg *et al*., 2012; Lareen *et al*., 2016), this system provides a stringent empirical framework for testing whether prevalence-based cores preserve interpretable ecological structure across fundamentally different community types while substantially reducing data complexity. Specifically, we evaluated information retention across multiple levels of ecological organization, including community composition, predicted functional profiles, and microbial association networks.

## Methods

### Experimental design

The study was conducted within the PhytOakmeter network (https://gepris.dfg.de/project/507084794), in which phytometers of the clonal pedunculate oak genotype *Quercus robur* DF159, generated via micro-propagation to retain their common genetic identity, were out-planted at 12 experimental sites distributed along a broad latitudinal gradient across France, Germany, and Finland (Herrmann *et al*., 2016). At each site, 12 clonal trees were planted within 40 × 40 m plots between 2013 and 2016, with planting dates varying among sites. All sites were equipped with environmental sensors recording air and soil temperature as well as soil moisture. Soil physicochemical properties were additionally characterized, revealing pronounced environmental heterogeneity across the transect (Table S1). This multi-site design provides a replicated system for studying plant-associated microbiomes across contrasting environments, allowing robust evaluation of whether ecological patterns remain consistent across compartments despite this pronounced spatial and climatic heterogeneity.

### Rhizosphere, roots, and leaves sampling

Between May and June 2024, six trees per site were sampled. For each tree, rhizosphere, root, and leaf compartments were collected to capture distinct plant-associated microbial habitats.

Rhizosphere and root samples were obtained by tracing lateral roots originating from the trunk. Soil was carefully excavated until at least two lateral roots per tree were exposed, in order to account for small-scale spatial heterogeneity. Fine roots with adhering soil were collected and transferred to sterile containers. Rhizosphere samples were defined as soil tightly associated with roots and influenced by root exudation (Vishwakarma *et al*., 2020). Adhering soil was gently removed by shaking and further separated from root material using sterile brushes. Only soil directly associated with root surfaces was retained, while loosely attached or falling soil particles, as well as visible debris (e.g., root fragments, plant material, stones), were manually removed to ensure compartment specificity (Barillot *et al*., 2013; Uroz *et al*., 2016). The resulting rhizosphere soil was immediately flash-frozen in liquid nitrogen. After removal of rhizosphere soil, brushed roots were collected separately and flash-frozen.

For the leaf compartment, two healthy, non-symptomatic leaves per tree were harvested from the Lammas shoot flush, i.e. the secondary burst of tree growth (Hilton *et al*., 1987), placed in sterile containers, and immediately flash-frozen in liquid nitrogen.

### Molecular sample processing and amplicon sequencing

DNA extraction and sequencing preparation followed the protocol described by (Habiyaremyeet al., 2020), with minor modifications for root and leaf samples. Prior to DNA extraction, root and leaf tissues were flash-frozen in liquid nitrogen and homogenized into a fine powder using a mortar and pestle. DNA was extracted using the PowerSoil DNA Isolation Kit (Qiagen, Hilden, Germany) following the manufacturer’s instructions. To ensure comparability across sample types, 120 mg of material was used for root and leaf samples, and 300 mg for rhizosphere samples. DNA was eluted in 60 μL (roots and leaves) or 100 μL (rhizosphere soil). All extracts were quantified using a NanoDrop spectrophotometer and normalized to 20 ng/μL prior downstream processing.

Bacterial and fungal community composition was assessed via amplicon sequencing. Bacterial communities were targeted using the V4 region of the 16S rRNA gene with primers P5-8N-515F + P5-7N-515F/P7-2N-806R + P7-1N-806R (Apprill *et al*., 2015; Parada *et al*., 2016). To reduce oak-derived contamination in leaf samples, a mitochondria- and chloroplast-specific blocking oligonucleotide was included during PCR amplification (Hussain *et al*., 2025). Fungal communities were targeted by amplifying the internal transcribed spacer 2 (ITS2) region using the P5-5N-ITS4 + P5-6N-ITS4/ P7-3N-fITS7 + P7-4N-fITS7 primer mix (Gardes & Bruns, 1993; Ihrmark *et al*., 2012; Leonhardt *et al*., 2019). All PCR amplifications were performed in triplicate under conditions described by Habiyaremye et al. (2020).

PCR products were pooled at equimolar concentrations, purified, and indexed using Nextera XT adaptors. After a second purification step, DNA libraries were normalized, pooled and quality-checked for fragment size distribution. Sequencing was performed on an Illumina MiSeq platform using a 2 × 300 bp paired-end chemistry at the Soil Ecology Department of the UFZ – Helmholtz Centre for Environmental Research in Halle (Saale), Germany.

### Bioinformatics

Raw sequencing data were processed using the dadasnake pipeline (v0.11.2; Weißbecker et al. 2021), which implements a workflow based on the DADA2 algorithm (Callahan *et al*., 2016). Default parameters were used unless otherwise specified.

Briefly, paired-end reads were quality-filtered based on Phred scores (Q13 for 16S and Q15 for ITS), and low-quality reads were removed. Sequencing errors were modeled and corrected using the DADA2 algorithm, and identical reads were collapsed into amplicon sequence variants (ASVs) as an intermediate, error-corrected sequence representation. Chimeric sequences were identified and removed using the DADA2 consensus approach. ASVs were subsequently clustered into operational taxonomic units (OTUs) using a 97% sequence similarity threshold (Estensmo *et al*., 2021; Schloss, 2021), which served as the final ecological unit for downstream community analyses. Taxonomic classification was performed by assigning sequences to taxa using the SILVA database (v138.1; (Quast *et al*., 2013)) for bacterial 16S sequences, and the UNITE database (v10.0; (Nilsson et al., 2019)) for fungal ITS2 sequences. Sequences with ambiguous or unclassified taxonomy were further examined using BLAST-based searches to improve taxonomic resolution.

### Data processing and statistical analysis

Bacterial and fungal OTU and taxonomy tables were processed in R (v4.3.1; R Core Team 2026) using the phyloseq package (v1.46.0; McMurdie & Holmes, 2013). OTUs not assigned at least to phylum level, as well as mitochondrial or chloroplast sequences, were removed prior to analyses.

To avoid arbitrary prevalence cutoffs, the prevalence threshold was determined using a data-driven, inflection-point-based approach (v1.3.6; Christopoulos 2025). We related OTU prevalence to the number of retained OTUs, fitted a smoothed spline, and applied the Extremum Surface Estimator (ESE) to identify the point at which the curve stabilizes, using a span of 0.9 based on a sensitivity analysis showing it reliably yields a valid, stable core threshold across all sample compartments (Fig. S1). ESE was preferred over the alternative Extremum Distance Estimator (EDE), which instead marks the onset of change in OTU loss, because our goal was to capture stabilization rather than its onset. This provided a reproducible, data-driven definition of the core community.

Rarefaction was performed after core identification, preserving the prevalence structure used for core inference while controlling for sequencing depth in downstream analyses; for comparisons between core and original datasets, the full datasets were rarefied accordingly.

To assess the robustness of the core definition under reduced sampling effort, samples were iteratively and randomly removed until only three remained, with each scenario repeated across 100 permutations to quantify variability in the resulting core threshold. Nonlinear trends between sample size and core stability were modelled using LOESS regression (span = 0.5).

To validate whether the identified core generalized beyond the dataset used to define it, we performed a leave-one-site-out (LOSO) cross-validation. For each study site group (n = 12), the core threshold was re-estimated using only the remaining eleven groups, applying the same prevalence-based core detection procedure described above. OTUs meeting this threshold were designated as the core set for that fold, and core recovery was calculated as the proportion of these OTUs detected in the held-out site, which had not contributed to core definition. This was repeated across all 12 folds for both the bacterial and fungal datasets and the three compartments. High and consistent recovery across folds would indicate that the core reflects a genuine pattern that generalizes across the sampled environmental gradient, rather than a reconstruction specific to the source dataset.

To evaluate whether the core preserved ecological structure, we compared Bray–Curtis dissimilarity matrices (aligned by shared samples) between original and core datasets using Mantel tests, as a measure of information retention after dimensionality reduction, and PERMANOVA, to test whether patterns of site differentiation were conserved after core filtering.

Functional potential of bacterial communities was inferred using PICRUSt2 (Douglas *et al*., 2020), and KEGG Orthology (KO) profiles (Kanehisa *et al*., 2016)) were compared between original and core datasets at the level of the full KO set (28,067 functions) and a subset of plant growth-promoting rhizobacteria (PGPR; 4,959 KOs; Table S2). Fungal lifestyles were analyzed analogously using a curated FungalTraits database (Põlme et al., 2021);Table S3). Mantel tests quantified similarity between functional profiles of original and core communities across compartments.

To test whether core membership was driven primarily by abundance rather than prevalence, OTU abundances were compared between retained and excluded taxa using ANOVA, and the distribution of the 100 most abundant OTUs was visualized to assess this relationship.

Microbial association networks were inferred from core datasets using SPIEC-EASI (Kurtz et al., 2015) with Meinshausen–Bühlmann neighborhood selection; non-rarefied data were used, as SPIEC-EASI internally applies a centered log-ratio transformation. Networks were built exclusively from core taxa, with no direct comparison to networks derived from the full (non-core) dataset; our aim was to assess whether the core alone yields stable, structurally coherent networks. The most stable network was selected across 20 lambda steps (5e-2 to 1e-4, 50 stability permutations) based on stability and structural coherence (connectivity, density, and the presence of a single dominant component). Network topology was characterized using degree, closeness, and betweenness centrality, with taxa in the top 5% of any of these three metrics classified as highly central taxa (putative network hubs).

## Results

### A prevalence-based core dramatically reduces community complexity

The data-driven prevalence threshold retained only a small fraction of the original microbial diversity across all compartments. After core identification and rarefaction, bacterial (16S) core communities contained less than 6% of the original OTUs, while fungal (ITS) cores conserved less than 3% (Table 1; Table S4). Despite this substantial reduction, sufficient sequencing depth was maintained for downstream analyses. Following rarefaction of the core dataset, only five samples were excluded, while no OTUs were lost (Table 1).

**Table 1.** Summary of the core microbiome filtering results for bacterial and fungal communities across leaf, roots, and rhizosphere compartments. Final OTUs, reads, and samples retained after filtering are reported together with the number of samples lost during the process.

| Compartment | Marker | Initial OTUs | Initial reads | Initial Samples | Prevalence threshold | Final OTUs | Final reads | Final Samples | Lost Samples | % conserved read | % conserved OTUs |
| --- | --- | --- | --- | --- | --- | --- | --- | --- | --- | --- | --- |
| Leaves | 16S | 3320 | 1405614 | 59 | 0.48484 | 64 | 103008 | 58 | 1 | 7.32832 | 1.92771 |
| Rhizosphere | 16S | 6093 | 3942068 | 70 | 0.69696 | 316 | 993510 | 70 | 0 | 25.20276 | 5.18627 |
| Roots | 16S | 5038 | 3094826 | 69 | 0.61616 | 312 | 982153 | 67 | 2 | 31.73532 | 6.19293 |
| Leaves | ITS | 1154 | 2686554 | 70 | 0.47474 | 34 | 261096 | 69 | 1 | 9.71862 | 2.94627 |
| Rhizosphere | ITS | 2988 | 2680137 | 70 | 0.48484 | 33 | 129360 | 70 | 0 | 4.82661 | 1.10441 |
| Roots | ITS | 2571 | 2506054 | 69 | 0.48484 | 24 | 51884 | 68 | 1 | 2.07034 | 0.93348 |

Core simulations revealed that prevalence thresholds were robust across sampling effort. Although threshold values increased when sample numbers became very small, they remained remarkably stable above approximately 15 samples (Fig. S2). Variability among simulations differed between compartments, with more complex bacterial root and rhizosphere communities showing greater variance than leaves, while fungal communities exhibited comparatively stable thresholds (Fig. S2; Table 1; Table S5). These analyses indicate that the identified cores represent robust subsets of the original communities rather than artifacts of sample size.

Core taxon recovery was high overall for both markers, and LOSO cross-validation confirmed these patterns held across held-out sites (Fig. 2; Table S6). For 16S, mean recovery was strong and consistent across all three compartments (∼92–93%, SD ∼10–13%; Table S6); the three compartments reached 100% recovery in at least one site, with even the lowest-recovering sites (56.7–69.6%; Fig. 2a-c, Table S7) still capturing the majority of the core. For ITS, recovery was also high overall, highest in leaves (93.8% ± 6.3%), followed by roots (88.7% ± 11.5%) and rhizosphere (83.6% ± 10.0%; Fig. 2d-f; Table S6). Several sites in roots and leaves reached full recovery. Across both markers, leaves showed the most consistent recovery of all compartments.

**Fig 2.**
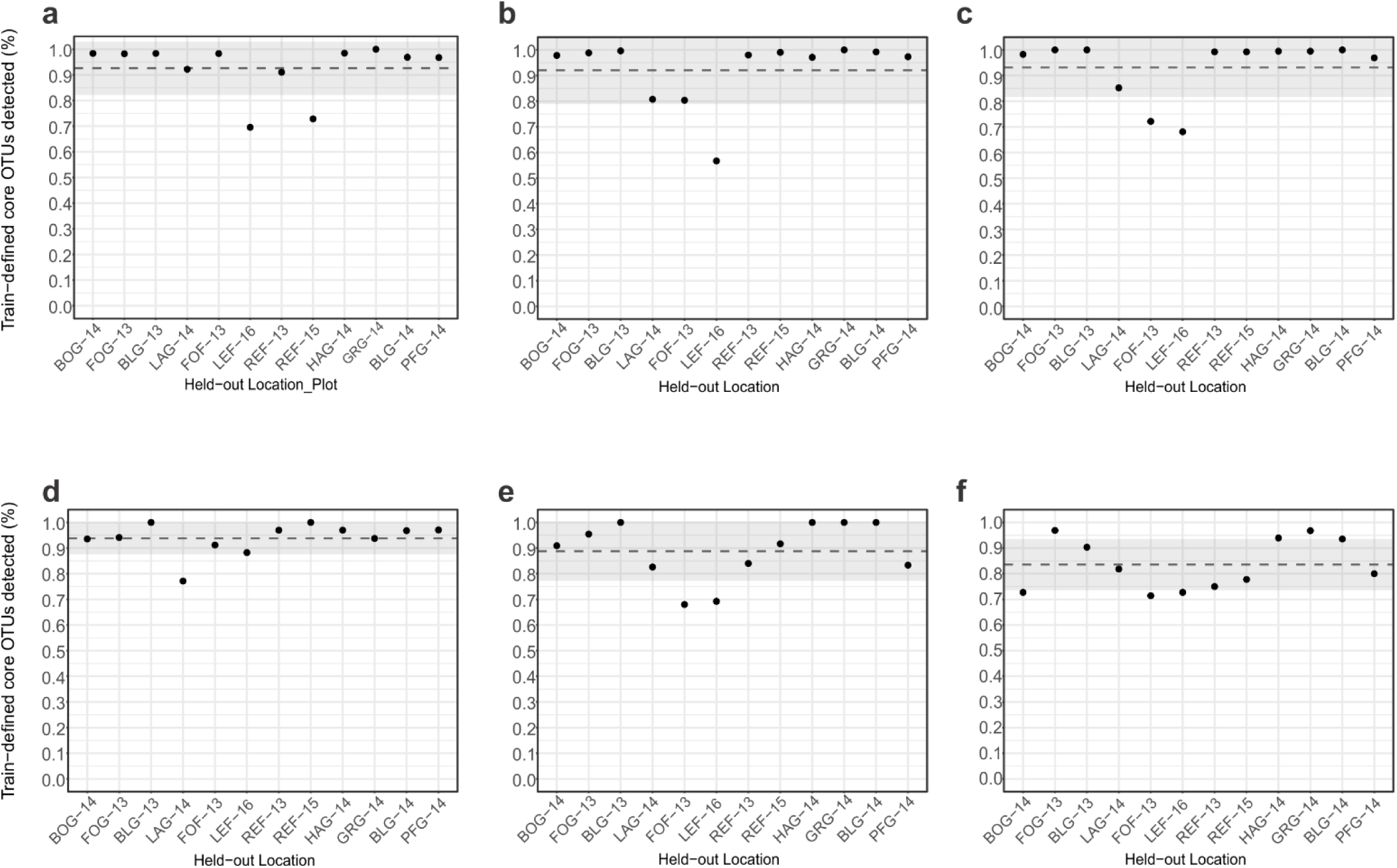
Leave-one-site-out (LOSO) validation of core microbiome recovery. Points show the fraction of train-defined core OTUs detected in each held-out site; dashed line and grey band show mean ± 1 SD across sites. (a) Bacterial leaves, (b) Bacterial roots, (c) Bacterial rhizosphere, (d) Fungal leaves, (e) Fungal roots and (f) Fungal rhizosphere.

### Core communities preserve the dominant ecological structure of microbial communities

Despite only representing a small fraction of the original diversity, core communities retained the major patterns of community composition.

Across bacterial communities, Mantel correlations between Bray-Curtis distance matrices of the original and core dataset were consistently high across compartments (r= 0.92-0.99), indicating that the reduced dataset retained nearly all multivariate relationships among the samples (Fig. 3a-c). Fungal communities showed lower, but still significant, correlations (r = 0.68–0.90), indicating that community simplification remained effective despite the greater heterogeneity of ITS datasets (Fig. 3d-e).

**Fig. 3.**
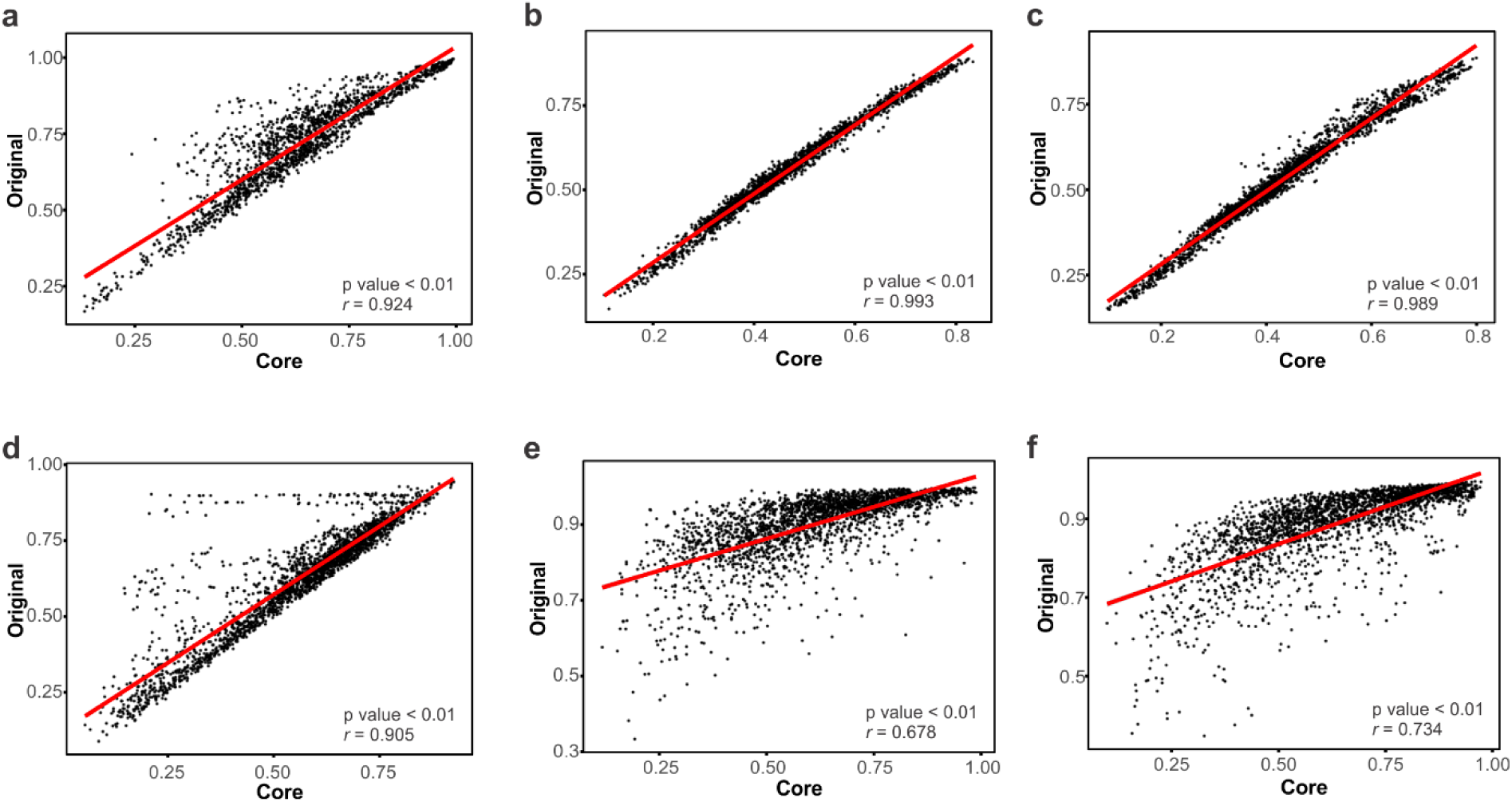
Mantel test correlation between the bray Curtis distance metric of the original and the core dataset. The top plots represents the correlations between 16S matrix distances for a) leaves, b) roots and c) rhizosphere and the lower plots for the ITS matrix distances for d) leaves, e) roots and f) rhizosphere.

Consistent with these results, PERMANOVA analyses produced highly concordant outcomes between original and core datasets across all compartments (Fig. S3; Table S8; Table S9). Most pairwise comparisons yielded identical statistical conclusions before and after dimensional reduction, particularly for bacterial communities. Discordant results were rare and occurred mainly within fungal datasets.

Together, these findings demonstrate that the vast majority of ecological information underlying community differentiation is preserved despite the removal of more than 94-97% of observed taxa.

### Functional information is largely retained after dimensional reduction

The strong conservation of community structure was accompanied by a similarly high retention of predicted functional information.

For bacterial communities, between 69% and 81% of predicted KEGG orthologs present in the original datasets were retained within the corresponding core communities (Table S10). Similar patterns were observed for plant growth-promoting functional categories. Across all compartments, functional profiles of original and core datasets were highly correlated (r = 0.88 - 0.94; Fig. S4a-c).

Fungal communities retained fewer functional traits than bacteria, reflecting their lower taxonomic richness and greater compositional heterogeneity. Nevertheless, significant correlations between original and core functional profiles were detected in all compartments, with the strongest relationship observed in leaves (Fig. S4d-f).

These results indicate that prevalence-based dimensional reduction conserves much of the functional information represented within microbial communities despite substantial taxonomic simplification.

### Core communities preferentially retain abundant taxa while excluding locally restricted taxa

Across all bacterial compartments, taxa retained within the core were significantly more abundant than taxa excluded during filtering (Fig. 4). However, prevalence and abundance were not equivalent. Several highly abundant OTUs were absent from the core because they occurred only in a limited number of samples. This pattern was particularly evident within fungal rhizosphere communities, where the most abundant OTU of the complete dataset was not retained in the core (Fig. S5).

**Fig. 4.**
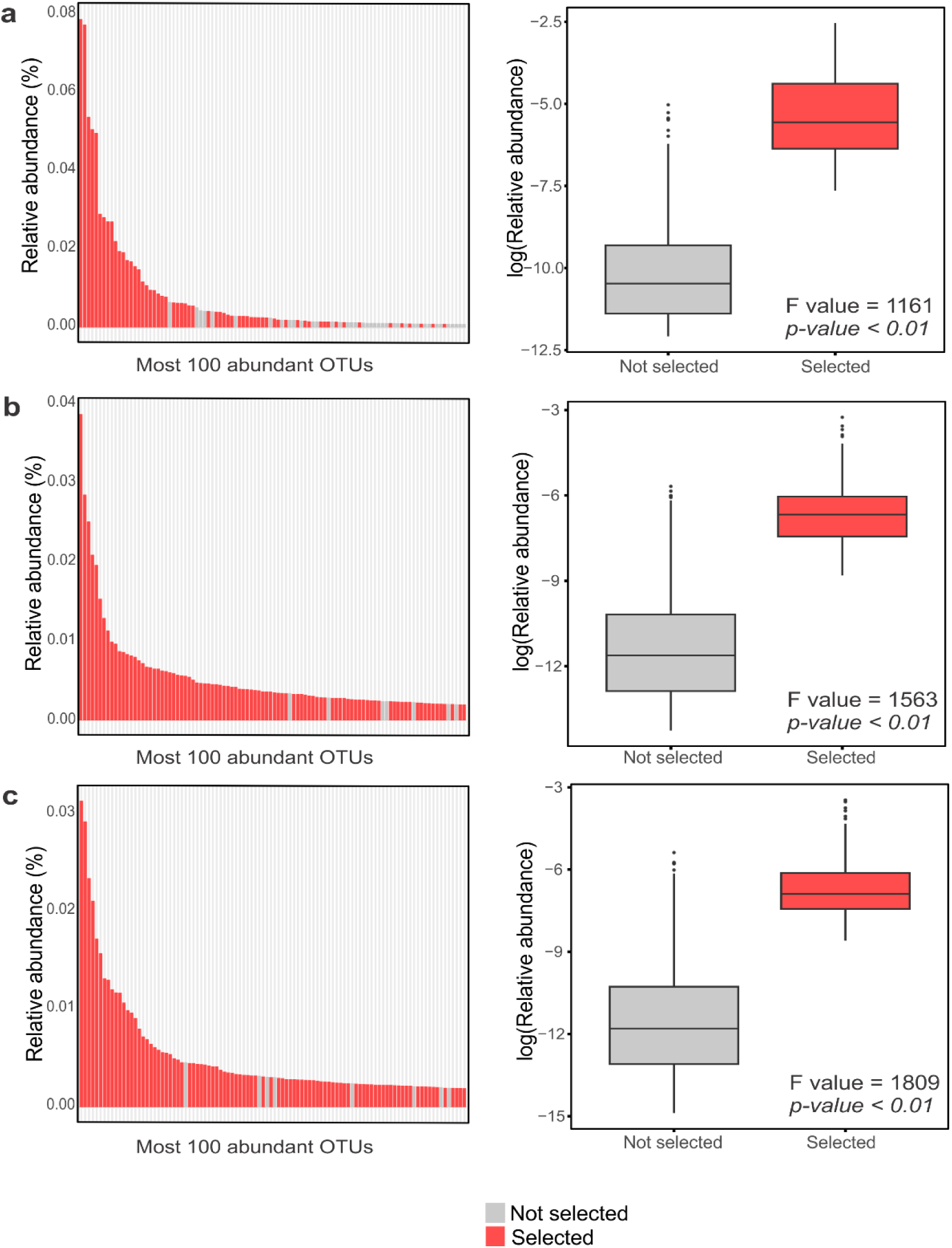
Relative abundance distribution of the 100 most abundant 16S OTUs across a) leaves, b) roots and c) rhizosphere. In the left panels, OTUs highlighted in red correspond to taxa selected by the core microbiome identification approach, whereas gray bars represent non-selected taxa. Right panels show boxplots comparing the log-transformed relative abundance of selected and non-selected OTUs.

These findings confirm that prevalence-based filtering preferentially captures ecologically consistent taxa rather than simply selecting the most abundant members of the community.

### Higher-order interaction structure is preserved in core-derived networks

To evaluate whether dimensional reduction also preserves higher-order ecological organization, microbial association networks were reconstructed for core communities using SPIEC-EASI (Table S11). Network stability analyses identified optimal regularization parameters across compartments and marker genes, confirming that inferred network structures were robust to methodological variation.

Despite strong reductions in taxonomic richness, core-derived networks retained their fundamental structural properties. Bacterial networks were highly connected in root and rhizosphere compartments, forming single dominant components, whereas leaf-associated networks were more fragmented (Fig. 5). Fungal networks showed consistently higher fragmentation across all compartments (Fig. S6), reflecting greater ecological heterogeneity compared to bacterial communities.

**Fig. 5.**
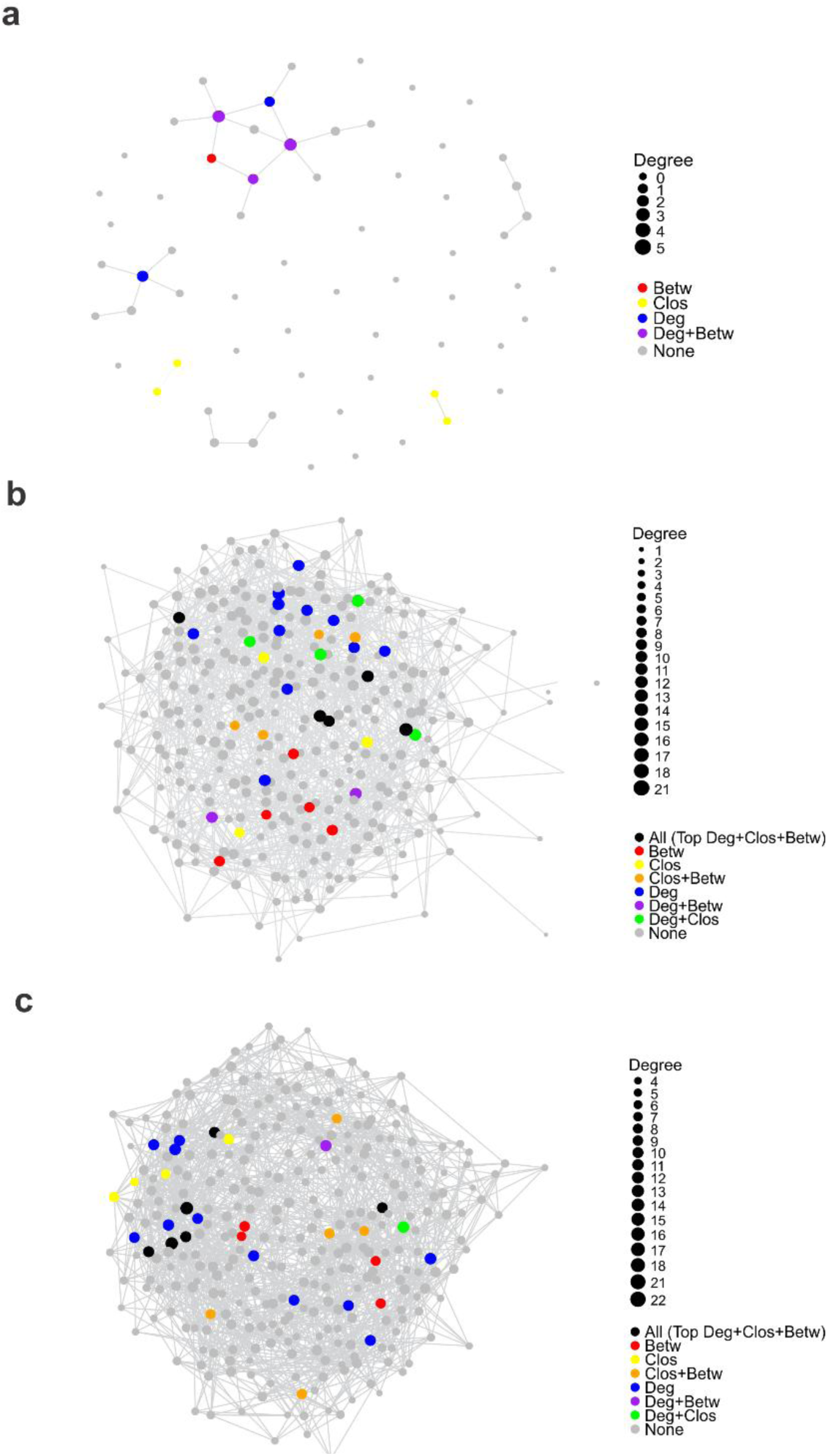
Network topology of the 16S microbial co-occurrence networks for (a) leaves, (b) roots, and (c) rhizosphere. Node size is proportional to degree (number of connections per node). Colored nodes highlight taxa identified as central according to different network centrality metrics: betweenness (Betw), closeness (Clos), degree (Deg). Gray nodes were not identified as central by any metric (“None”). Edges represent significant associations between taxa.

Across all datasets, central taxa could be identified within core networks based on multiple centrality metrics, although overlap between metrics was limited (Fig. 5, Fig. S5, Table S12). Overall, these results indicate that bacterial communities form highly interconnected networks with a single component, whereas fungal communities are more fragmented, likely due to ecological heterogeneity. This analysis highlights the OTUs that are most central within each compartment, providing insight into potential key taxa driving microbial interactions in leaves, roots, and rhizosphere.

## Discussion

Understanding how microbial communities can be meaningfully simplified without losing essential ecological information remains a central challenge in microbial ecology. In this study, we demonstrate that a prevalence-based, data-driven core microbiome definition can substantially lower the diversity of the filtered microorganisms while preserving key aspects of community structure, function, and higher-order interaction patterns across multiple plant-associated compartments along a wide environmental gradient.

Across all datasets, core microbial communities retained only a small fraction of total diversity (<6% of bacterial OTUs and <3% of fungal OTUs), yet preserved the dominant ecological structure of the original communities. Strong concordance in beta-diversity patterns between original and core datasets indicated that major gradients in community composition remained largely intact despite the extreme dimensional reduction. This suggests that much of the multivariate ecological signal in microbial communities is concentrated within a relatively small subset of consistently occurring taxa.

Importantly, this reduction is not equivalent to a loss of ecological information, but rather reflects a filtering of stochastic or locally restricted variations (Bissett *et al*., 2010; Bokulich *et al*., 2014; Gourmelon *et al*., 2016). Prevalence-based selection inherently emphasizes taxa that are consistently detected across samples, thereby capturing stable ecological signal while excluding transient or spatially constrained taxa (Shapira, 2016; Risely *et al*., 2021). This mechanism explains why, despite pronounced variation in climate, geography, and local edaphic conditions, core communities retained strong compositional and functional coherence even with drastic reductions in taxonomic richness (Delgado-Baquerizo *et al*., 2018; Toju *et al*., 2018). Notably, Lee et al. (2026), point out that core definitions often implicitly conflate two distinct objectives: identifying recurrent membership and optimizing the explanatory reconstruction of a dataset. Our approach explicitly targets the former while retaining strong explanatory power: it identifies the most consistent OTUs in the dataset without losing the identity and differences among communities.

The preservation of functional profiles further supports this interpretation. Although the core represented a small fraction of taxa, it retained a substantial proportion of predicted functional potential, including plant growth-promoting traits, indicating that functional structure is partially decoupled from taxonomic richness and is concentrated in a subset of widely distributed taxa (Delgado-Baquerizo *et al*., 2018; Louca *et al*., 2018). This supports the ecological relevance of the core as a reduced representation of complex microbial communities (Shade & Handelsman, 2012), and indicates that the taxa retained within it still play fundamental ecological roles despite representing only a small fraction of total diversity. Different OTUs may perform similar ecological functions in different samples and sites despite differing in their taxonomic identity (Cohan, 2002). As a result, functionally redundant OTUs can occur in different subsets of samples while maintaining equivalent ecosystem functions, leading to numerous structural zeros (Kaul *et al*., 2017) in OTU abundance matrices. Under these conditions, a core based on functional similarity is a fundamentally different criterion from a prevalence-based core microbiome, because shared function does not imply the consistent occurrence of the same OTUs across samples. However, it is important to consider that our results should be interpreted as potential functional capacity inferred from marker-gene data rather than direct measurements of microbial activity (Douglas *et al*., 2020): a limitation inherent to amplicon-based functional prediction in general, not specific to prevalence-based core definitions. This distinction matters for how the functional convergence criterion discussed below should be interpreted: our results demonstrate that predicted functional potential is retained within the core, which is a necessary but not sufficient condition for claiming conserved biological function.

A key conceptual outcome of this study is that abundance and prevalence represent distinct ecological dimensions that should not be conflated. Highly abundant taxa were not necessarily part of the core microbiome if their occurrence is spatially restricted (Custer *et al*., 2023),while less abundant taxa were consistently retained due to their broad distribution across samples (Cordero & Polz, 2014; Louca *et al*., 2018). This highlights that prevalence-based cores capture ecological consistency rather than dominance, reinforcing the idea that ecological importance cannot be inferred from abundance alone (Lynch & Neufeld, 2015; Jousset *et al*., 2017). This result is consistent with Bray-Curtis-based and abundance-occupancy approaches (Shade & Stopnisek, 2019; Lee *et al*., 2026), reinforcing that our prevalence-based cores also reflect the origin and identity of the samples.

Differences between bacterial and fungal communities further emphasize this point. Fungal communities exhibited lower structural and functional retention than bacterial communities, reflecting their higher spatial heterogeneity and stronger environmental filtering (Egidi *et al*., 2019). This suggests that prevalence-based reduction is more effective in systems with strong community coherence, whereas more heterogeneous systems may contain a larger proportion of context-dependent taxa that are filtered out under core definitions (O’Brien *et al*., 2005; Tedersoo *et al*., 2014).

At a level of higher-order organization, networks reconstructed from core communities using SPIEC-EASI showed biologically plausible and stable topology across compartments and marker genes (Berry & Widder, 2014; Röttjers & Faust, 2018). Bacterial networks were highly connected in root and rhizosphere compartments, forming single dominant components, whereas leaf-associated and fungal networks were more fragmented (Xiong *et al*., 2021)(Sun *et al*., 2017), likely reflecting greater ecological heterogeneity in these systems.

Simulation analysis further validates the prevalence-based core as a reliable and adaptive framework: threshold behavior was driven by intrinsic community complexity rather than sampling artifacts. Thresholds increased only at very low sample sizes, stabilizing once a baseline sampling depth was reached, i.e., rapidly in fungal groups and bacterial leaves, more gradually in root and rhizosphere bacteria, consistent with the higher richness and larger rare biosphere of these belowground compartments (Baldrian, 2019)(Elshahed *et al*., 2008; Lynch & Neufeld, 2015; Hermans *et al*., 2019). LOSO cross-validation confirmed that defined cores generalize well to sites that did not contribute to their definition, ruling out overfitting to the full dataset. The stronger and more consistent recovery of the bacterial core relative to the fungal core aligns with broader evidence that fungal community assembly is more strongly shaped by dispersal limitation, while bacterial assembly is more structured by local environmental filtering (Chen *et al*., 2020; Zhang *et al*., 2021), which explains the lower and more variable ITS recovery, particularly belowground.

This interpretation also helps reconcile previously conflicting definitions of the core microbiome. Function-based, abundance-based, and prevalence-based approaches often identify partially non-overlapping sets of taxa(Risely, 2020). Our results suggest this arises because each criterion captures a different aspect of community structure, i.e., prevalence-based cores specifically retain the most stable component of ecological signal, whereas other definitions emphasize dominance or functional potential. This aligns with Lee et al. (2026), who call for greater terminological precision and propose restricting “core microbiome” to taxa explicitly shown to be conserved across spatial, temporal, and environmental dimensions and linked to conserved ecological function, while taxa selected primarily for their explanatory value should instead be termed “explanatory subsets of taxa.” Measured against this stricter standard, our core meets the spatial and environmental criteria robustly (12 sites, LOSO cross-validation) and shows strong explanatory power, but, as noted above, its functional support remains predictive rather than directly measured, and, as discussed below, temporal conservation was not assessed in the present design. Rather than treating this as a shortfall of our framework, we see it as clarifying what a prevalence-based core does and does not demonstrate. Consistent with this call for precision, rather than viewing the core as a biologically meaningful subset of dominant or “important” taxa, we propose understanding it as a data-driven, information-preserving abstraction. This abstraction reduces the zero-inflation of the original matrix while capturing the stable multivariate structure of microbial communities and substantially reducing the dimensionality and complexity of the dataset. This framing is also intended to address the major limitations of common metabarcoding data (Gold *et al*., 2023), while intentionally remaining open to other approaches. We do not claim this prevalence-based approach is the only valid solution; approaches such as Bray-Curtis-based selection (Shade & Stopnisek, 2019; Lee *et al*., 2026), which we found broadly consistent with our results above, represent one promising alternative among several, and, like ours, remain to be tested with the same degree of systematic robustness scrutiny across multiple sites, compartments, and markers.

Finally, the flexibility of our framework allows core definitions to be adapted to different spatial or temporal scales, making prevalence-based core identification a useful tool for simplifying complex microbial datasets while maintaining ecological interpretability. Because sampling was conducted within a single growing season, our study cannot directly evaluate temporal stability, one of the criteria emphasized by Lee et al. (2026), and this remains an important next step, ideally through repeated sampling within the same long-term experimental network (e.g., PhytOakmeter) across multiple years, which would allow the core to be tested not only across space but also across time within an already-established, environmentally characterized system, and would move it closer to satisfying the full set of criteria for a biologically conserved core microbiome rather than a reproducible explanatory representation. Future work should also explore how disturbance and host identity influence the stability of these information-preserving cores, and whether shifts in community structure are primarily reflected in core membership or in the surrounding variable microbial pool.

## Supporting information

Overview of experimental study sites included in this study.

KEGG Orthology (KO) terms identified per compartment and plant growth-promoting rhizobacteria (PGPR)-associated genes.

FUNGuild functional traits assigned per compartment.

Core microbiome OTUs with relative abundance per compartment, family, and genus, for bacteria and fungi.

Sample-subsampling simulation results for core microbiome definition across compartments and markers.

Summary of core taxon recovery by marker and plant compartment from LOSO analysis.

Leave-one-site-out (LOSO) cross-validation results for core microbiome definition across compartments and markers.

Summary of PERMANOVA significance consistency between original and core datasets across compartments and marker genes.

Pairwise PERMANOVA results comparing community composition between sites within each compartment and marker.

Functional overlap between the complete KEGG Orthology (KO) database and the core microbiome across leaf, root, and rhizosphere compartments.

SPIECEASI Network topology and stability metrics simulations of microbial networks inferred for bacterial and fungal core microbiomes across compartme

Co-occurrence network properties of core OTUs per compartment and marker, inferred using SPIECEASI.

## Acknowledgements

This work was carried out within the framework of “PhytOakmeter”, a DFG-funded Research Unit (project grant 507084794, individual subproject funding PR 2235/3-1 and GO 3507/5-1) and the Swiss National Science Foundation (SNSF). Our biggest gratitude belongs to François Buscot and Sylvie Herrmann as the intellectual founders of the PhytOakmeter. Moreover, we thank Lars Opgenoorth, Marie-Lara Bouffaud, Benjamin Dauphin, Martin Gossner, Katrin Heer, Mika Tarkka, Julia Baumeister, and Jasmina Rakovic for project coordination and administration. We thank the Helmholtz Centre for Environmental Research (UFZ) for logistic support and permission to do research on their sites. We further acknowledge local support at each site: Dr. Ines Merbach (Experimental Field Station Bad Lauchstädt), Julien Parmentier and Marie-Laure Greil (INRAE – Unité Expérimentale Arboricole – Toulenne), Xavier Lacroix (Commune de Fontain), Markku Rantala (Natural Resources Institute Finland, Luke), Prof. Ernst-Detlef Schulze (Forest of Rehungen), and Andreas Sickert (Urban Forestry of Leipzig), as well as the UFZ Terrestrial Environmental Observatories (TERENO). We further thank Ines Krieg, Barbara Krause and Stefan Tautkus for producing the DF159 oak clones, maintaining and monitoring of these sites. We thank Melanie Günther and Beatrix Schnabel for their help in the wet lab. The sequencing data was analyzed at the High-Performance Computing Cluster EVE, a joint effort between the UFZ and the German Centre for Integrative Biodiversity Research (iDiv) Halle-Jena-Leipzig. To ensure data FAIRness, we utilized the DataPLANT personal assistance network and its tool stack, centered around the DataHUB.

## Author contributions

Conceptualization: all authors; Methodology: CQ; Software: CQ; Formal analysis: CQ; Investigation: all authors; Data curation: CQ, LP; Writing – original draft: CQ, KG; Writing – review & editing: all authors; Visualization: CQ; Supervision: LP, KG; Project administration: LP, KG; Funding acquisition: LP, KG.

## Data availability

To ensure data FAIRness, we utilized the DataPLANT personal assistance network and its tool stack, centered around the DataHUB. Our data publication and the code used can be found at at https://git.nfdi4plants.org/camilo-andres.quiroga-gonzalez/SP4a_European_Transect_Core, and the sequences used were stored at NCBI under BioProject accession number PRJNA1499632.

## References

Ainsworth TD, Krause L, Bridge T, Torda G, Raina JB, Zakrzewski M, Gates RD, Padilla-Gamiño JL, Spalding HL, Smith C, et al. 2015. The coral core microbiome identifies rare bacterial taxa as ubiquitous endosymbionts. ISME Journal 9: 2261–2274.

Apprill A, Mcnally S, Parsons R, Weber L. 2015. Minor revision to V4 region SSU rRNA 806R gene primer greatly increases detection of SAR11 bacterioplankton. Aquatic Microbial Ecology 75: 129–137.

Bäcker M, Doekes HM, Garza DR, Meijer J, van Vliet S, Allen RJ, Hogeweg P, Dutilh BE, van Dijk B. 2026. Spatial structure: shaping the ecology and evolution of microbial communities. FEMS microbiology reviews 50: fuaf067.

Bais HP, Weir TL, Perry LG, Gilroy S, Vivanco JM. 2006. The role of root exudates in rhizosphere interactions with plants and other organisms. Annual Review of Plant Biology 57: 233–266.

Baldrian P. 2019. The known and the unknown in soil microbial ecology. FEMS Microbiology Ecology 95: fiz005.

Banerjee S, Schlaeppi K, van der Heijden MGA. 2018. Keystone taxa as drivers of microbiome structure and functioning. Nature Reviews Microbiology 16: 567–576.

Barillot CDC, Sarde CO, Bert V, Tarnaud E, Cochet N. 2013. A standardized method for the sampling of rhizosphere and rhizoplan soil bacteria associated to a herbaceous root system. Annals of Microbiology 63: 471–476.

Berry D, Widder S. 2014. Deciphering microbial interactions and detecting keystone species with co-occurrence networks. Frontiers in Microbiology 5: 219.

Bissett A, Richardson AE, Baker G, Wakelin S, Thrall PH. 2010. Life history determines biogeographical patterns of soil bacterial communities over multiple spatial scales. Molecular Ecology 19: 4315–4327.

Bokulich NA, Thorngate JH, Richardson PM, Mills DA. 2014. Microbial biogeography of wine grapes is conditioned by cultivar, vintage, and climate. Proceedings of the National Academy of Sciences of the United States of America 111: E139–E148.

Callahan BJ, McMurdie PJ, Rosen MJ, Han AW, Johnson AJA, Holmes SP. 2016. DADA2: High-resolution sample inference from Illumina amplicon data. Nature Methods 13: 581–583.

Chakraborty D, Jousset A, Wei Z, Banerjee S. 2025. Rare taxa in the core microbiome. Trends in Microbiology 33: 727–737.

Chen J, Wang P, Wang C, Wang X, Miao L, Liu S, Yuan Q, Sun S. 2020. Fungal community demonstrates stronger dispersal limitation and less network connectivity than bacterial community in sediments along a large river. Environmental Microbiology 22: 832–849.

Christopoulos C. 2025. inflection: Finds the Inflection Point of a Curve.

Cohan FM. 2002. What are bacterial species? Annual Review of Microbiology 56: 457– 487.

Cordero OX, Polz MF. 2014. Explaining microbial genomic diversity in light of evolutionary ecology. Nature Reviews Microbiology 12: 263–273.

Custer GF, Gans M, van Diepen LTA, Dini-Andreote F, Buerkle CA. 2023. Comparative Analysis of Core Microbiome Assignments: Implications for Ecological Synthesis. mSystems 8: e01066–22.

Delgado-Baquerizo M, Oliverio AM, Brewer TE, Benavent-González A, Eldridge DJ, Bardgett RD, Maestre FT, Singh BK, Fierer N. 2018. A global atlas of the dominant bacteria found in soil. Science 359(6373): 320–325.

Douglas GM, Maffei VJ, Zaneveld JR, Yurgel SN, Brown JR, Taylor CM, Huttenhower C, Langille MGI. 2020. PICRUSt2 for prediction of metagenome functions. Nature Biotechnology 38: 685–688.

Egidi E, Delgado-Baquerizo M, Plett JM, Wang J, Eldridge DJ, Bardgett RD, Maestre FT, Singh BK. 2019. A few Ascomycota taxa dominate soil fungal communities worldwide. Nature Communications 10: 2369.

Elshahed MS, Youssef NH, Spain AM, Sheik C, Najar FZ, Sukharnikov LO, Roe BA, Davis JP, Schloss PD, Bailey VL, et al. 2008. Novelty and uniqueness patterns of rare members of the soil biosphere. Applied and Environmental Microbiology 74: 5422– 5428.

Estensmo ELF, Maurice S, Morgado L, Martin-Sanchez PM, Skrede I, Kauserud H. 2021. The influence of intraspecific sequence variation during DNA metabarcoding: A case study of eleven fungal species. Molecular Ecology Resources 21: 1141–1148.

Gardes M, Bruns TD. 1993. ITS primers with enhanced specificity for basidiomycetes - application to the identification of mycorrhizae and rusts. Molecular Ecology 2: 113– 118.

Gibbons SM, Gilbert JA. 2015. Microbial diversity-exploration of natural ecosystems and microbiomes. Current Opinion in Genetics and Development 35: 66–72.

Gold Z, Shelton AO, Casendino HR, Duprey J, Gallego R, Van Cise A, Fisher M, Jensen AJ, D’Agnese E, Allan EA, et al. 2023. Signal and noise in metabarcoding data. PLoS ONE 18.

Gourmelon V, Maggia L, Powell JR, Gigante S, Hortal S, Gueunier C, Letellier K, Carriconde F. 2016. Environmental and geographical factors structure soil microbial diversity in new caledonian ultramafic substrates: A metagenomic approach. PLoS ONE 11.

Habiyaremye J de D, Goldmann K, Reitz T, Herrmann S, Buscot F. 2020. Tree Root Zone Microbiome: Exploring the Magnitude of Environmental Conditions and Host Tree Impact. Frontiers in Microbiology 11: 749.

Hamady M, Knight R. 2009. Microbial community profiling for human microbiome projects: Tools, techniques, and challenges. Genome Research 19: 1141–1152.

Hanson CA, Fuhrman JA, Horner-Devine MC, Martiny JBH. 2012. Beyond biogeographic patterns: Processes shaping the microbial landscape. Nature Reviews Microbiology 10: 497–506.

Hermans SM, Buckley HL, Lear G. 2019. Perspectives on the Impact of Sampling Design and Intensity on Soil Microbial Diversity Estimates. Frontiers in Microbiology 10: 1820.

Herrmann S, Grams TEE, Tarkka MT, Angay O, Bacht M, Bönn M, Feldhahn L, Graf M, Kurth F, Maboreke H, et al. 2016. Endogenous rhythmic growth, a trait suitable for the study of interplays between multitrophic interactions and tree development. *Perspectives in Plant Ecology*, Evolution and Systematics 19: 40–48.

Hilton GM, Packham JR, Willis AJ. 1987. Effects of experimental defoliation on a population of pedunculate oak (Quercus robur L.). New Phytologist 107: 603–612.

Huse SM, Ye Y, Zhou Y, Fodor AA. 2012. A core human microbiome as viewed through 16S rRNA sequence clusters. PLoS ONE 7: e34242.

Hussain U, Downie J, Ellison A, Denman S, McDonald J, Cambon MC. 2025. Peptide nucleic acid (PNA) clamps reduce amplification of host chloroplast and mitochondria rRNA gene sequences and increase detected diversity in 16S rRNA gene profiling analysis of oak-associated microbiota. Environmental Microbiome 20: 14.

Ihrmark K, Bödeker ITM, Cruz-Martinez K, Friberg H, Kubartova A, Schenck J, Strid Y, Stenlid J, Brandström-Durling M, Clemmensen KE, et al. 2012. New primers to amplify the fungal ITS2 region - evaluation by 454-sequencing of artificial and natural communities. FEMS Microbiology Ecology 82: 666–677.

Jousset A, Bienhold C, Chatzinotas A, Gallien L, Gobet A, Kurm V, Küsel K, Rillig MC, Rivett DW, Salles JF, et al. 2017. Where less may be more: How the rare biosphere pulls ecosystems strings. ISME Journal 11: 853–862.

Kanehisa M, Sato Y, Kawashima M, Furumichi M, Tanabe M. 2016. KEGG as a reference resource for gene and protein annotation. Nucleic Acids Research 44: D457– D462.

Kaul A, Davidov O, Peddada SD. 2017. Structural zeros in high-dimensional data with applications to microbiome studies. Biostatistics 18: 422–433.

Knight R, Vrbanac A, Taylor BC, Aksenov A, Callewaert C, Debelius J, Gonzalez A, Kosciolek T, McCall LI, McDonald D, et al. 2018. Best practices for analysing microbiomes. Nature Reviews Microbiology 16: 410–422.

Lareen A, Burton F, Schäfer P. 2016. Plant root-microbe communication in shaping root microbiomes. Plant Molecular Biology 90: 575–587.

Lee J, Aponte Rolón B, de Lorimier P, Ané J, Benucci GMN, Carrell AA, Chai YN, Chandrasoma J, Geerdes N, Infante V, et al. 2026. Rethinking the soil core microbiome. New Phytologist.

Lemanceau P, Blouin M, Muller D, Moënne-Loccoz Y. 2017. Let the Core Microbiota Be Functional. Trends in Plant Science 22: 583–595.

Leonhardt S, Hoppe B, Stengel E, Noll L, Moll J, Bässler C, Dahl A, Buscot F, Hofrichter M, Kellner H. 2019. Molecular fungal community and its decomposition activity in sapwood and heartwood of 13 temperate European tree species. PLoS ONE 14: e0212120.

Louca S, Polz MF, Mazel F, Albright MBN, Huber JA, O’connor MI, Ackermann M, Hahn AS, Srivastava DS, Crowe SA, et al. 2018. Function and functional redundancy in microbial systems. Nature ecology & evolution, 2(6): 936–943.

Lundberg DS, Lebeis SL, Paredes SH, Yourstone S, Gehring J, Malfatti S, Tremblay J, Engelbrektson A, Kunin V, Rio TG Del, et al. 2012. Defining the core Arabidopsis thaliana root microbiome. Nature 488: 86–90.

Lynch MDJ, Neufeld JD. 2015. Ecology and exploration of the rare biosphere. Nature Reviews Microbiology 13: 217–229.

McMurdie PJ, Holmes S. 2013. Phyloseq: An R Package for Reproducible Interactive Analysis and Graphics of Microbiome Census Data. PLoS ONE 8: e61217.

Neu AT, Allen EE, Roy K. 2021. Defining and quantifying the core microbiome: Challenges and prospects. Proceedings of the National Academy of Sciences 118: e2104429118.

Nilsson RH, Larsson KH, Taylor AFS, Bengtsson-Palme J, Jeppesen TS, Schigel D, Kennedy P, Picard K, Glöckner FO, Tedersoo L, et al. 2019. The UNITE database for molecular identification of fungi: Handling dark taxa and parallel taxonomic classifications. Nucleic Acids Research 47: D259–D264.

O’Brien HE, Parrent JL, Jackson JA, Moncalvo JM, Vilgalys R. 2005. Fungal community analysis by large-scale sequencing of environmental samples. Applied and Environmental Microbiology 71: 5544–5550.

Parada AE, Needham DM, Fuhrman JA. 2016. Every base matters: Assessing small subunit rRNA primers for marine microbiomes with mock communities, time series and global field samples. Environmental Microbiology 18: 1403–1414.

Põlme S, Abarenkov K, Henrik Nilsson R, Lindahl BD, Clemmensen KE, Kauserud H, Nguyen N, Kjøller R, Bates ST, Baldrian P, et al. 2021. Correction to: FungalTraits: a user friendly traits database of fungi and fungus-like stramenopiles (Fungal Diversity, (2020), 105, 1, (1-16), 10.1007/s13225-020-00466-2). Fungal Diversity 107: 129–132.

Quast C, Pruesse E, Yilmaz P, Gerken J, Schweer T, Yarza P, Peplies J, Glöckner FO. 2013. The SILVA ribosomal RNA gene database project: Improved data processing and web-based tools. Nucleic Acids Research 41: D590–D596.

R Core Team. 2026. R: A Language and Environment for Statistical Computing.

Risely A. 2020. Applying the core microbiome to understand host–microbe systems. Journal of Animal Ecology 89: 1549–1558.

Risely A, Gillingham MAF, Béchet A, Brändel S, Heni AC, Heurich M, Menke S, Manser MB, Tschapka M, Wasimuddin M, et al. 2021. Phylogeny- and Abundance-Based Metrics Allow for the Consistent Comparison of Core Gut Microbiome Diversity Indices Across Host Species. Frontiers in Microbiology 12: 659918.

Röttjers L, Faust K. 2018. From hairballs to hypotheses–biological insights from microbial networks. FEMS Microbiology Reviews 42: 761–780.

Schloss PD. 2021. Amplicon Sequence Variants Artificially Split Bacterial Genomes into Separate Clusters. mSphere 6: 10–1128.

Shade A, Handelsman J. 2012. Beyond the Venn diagram: The hunt for a core microbiome. Environmental Microbiology 14: 4–12.

Shade A, Stopnisek N. 2019. Abundance-occupancy distributions to prioritize plant core microbiome membership. Current Opinion in Microbiology 49: 50–58.

Shapira M. 2016. Gut Microbiotas and Host Evolution: Scaling Up Symbiosis. Trends in Ecology and Evolution 31: 539–549.

Sun S, Li S, Avera BN, Strahm BD, Badgley BD. 2017. Soil bacterial and fungal communities show distinct recovery patterns during forest ecosystem restoration. Applied and Environmental Microbiology 83: e00966–17.

Tedersoo L, Bahram M, Põlme S, Kõljalg U, Yorou NS, Wijesundera R, Ruiz LV, Vasco-Palacios AM, Thu PQ, Suija A, et al. 2014. Global diversity and geography of soil fungi. Science 346: 1256688.

Toju H, Peay KG, Yamamichi M, Narisawa K, Hiruma K, Naito K, Fukuda S, Ushio M, Nakaoka S, Onoda Y, et al. 2018. Core microbiomes for sustainable agroecosystems. Nature Plants 4: 247–257.

Turnbaugh PJ, Ley RE, Hamady M, Fraser-Liggett CM, Knight R, Gordon JI. 2007. The Human Microbiome Project. Nature 449: 804–810.

Uroz S, Oger P, Tisserand E, CéBron A, Turpault MP, Bueé M, De Boer W, Leveau JHJ, Frey-Klett P. 2016. Specific impacts of beech and Norway spruce on the structure and diversity of the rhizosphere and soil microbial communities. Scientific Reports 6: 27756.

Vishwakarma K, Kumar N, Shandilya C, Mohapatra S, Bhayana S, Varma A. 2020. Revisiting Plant–Microbe Interactions and Microbial Consortia Application for Enhancing Sustainable Agriculture: A Review. Frontiers in Microbiology 11: 560406.

Weißbecker C, Schnabel B, Heintz-Buschart A. 2021. Dadasnake, a snakemake implementation of DADA2 to process amplicon sequencing data for microbial ecology. GigaScience 9: giaa135.

Xiong C, Zhu YG, Wang JT, Singh B, Han LL, Shen JP, Li PP, Wang GB, Wu CF, Ge AH, et al. 2021. Host selection shapes crop microbiome assembly and network complexity. New Phytologist 229: 1091–1104.

Zhang G, Wei G, Wei F, Chen Z, He M, Jiao S, Wang Y, Dong L, Chen S. 2021. Dispersal Limitation Plays Stronger Role in the Community Assembly of Fungi Relative to Bacteria in Rhizosphere Across the Arable Area of Medicinal Plant. Frontiers in Microbiology 12: 713523.

