## Supplementary Figures for "The core microbiome as a reproducible, data-driven abstraction, not a biological entity"

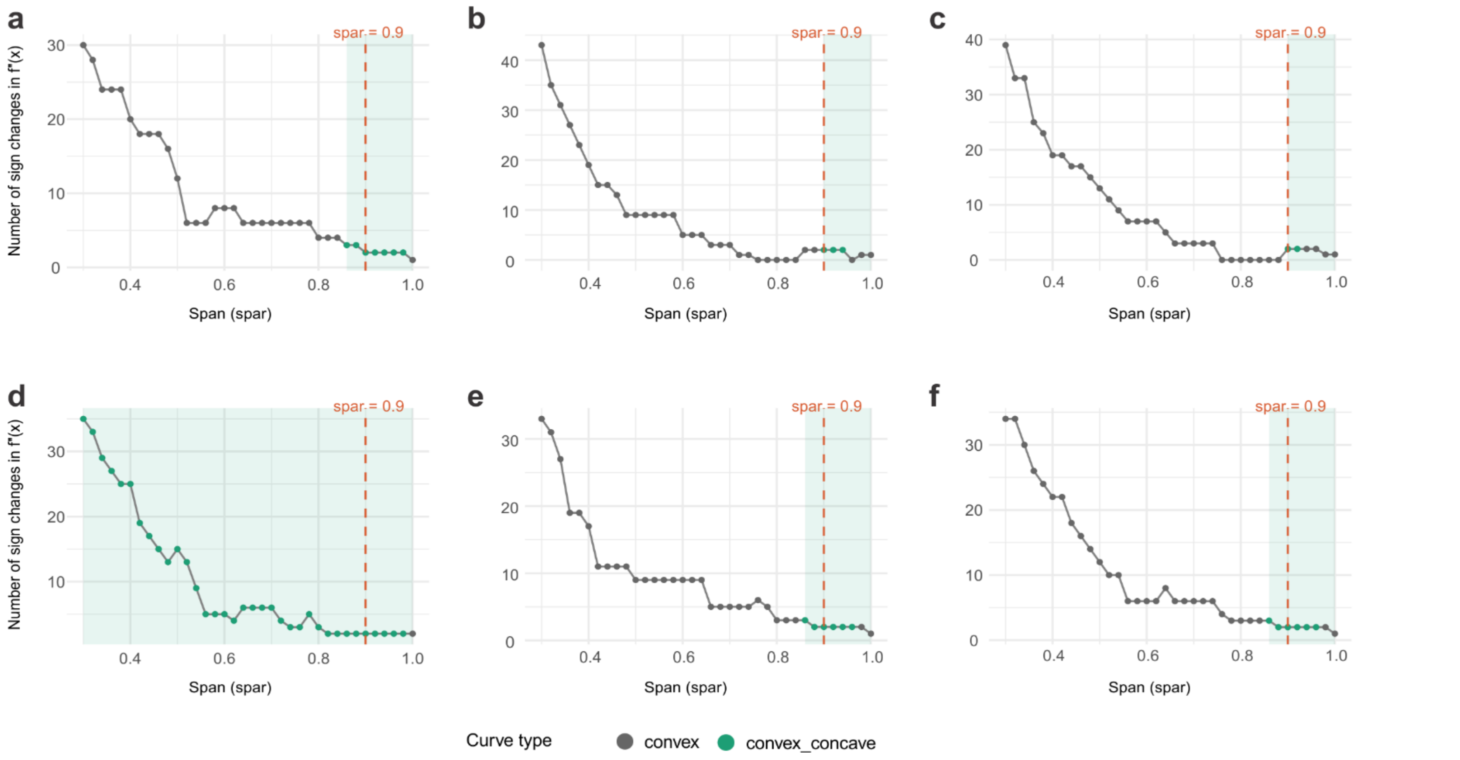


**Fig S1.** Sensitivity of ESE inflection detection to spline span (spar) across the six compartments. Grey = convex curves (no valid inflection); green = convex_concave curves satisfying the ESE assumption. Spar = 0.9 (dashed line) falls within the valid region in all panels: (a) leaves 16S, (b) roots 16S, (c) rhizosphere 16S, (d) leaves ITS, (e) roots ITS, (f) rhizosphere ITS.

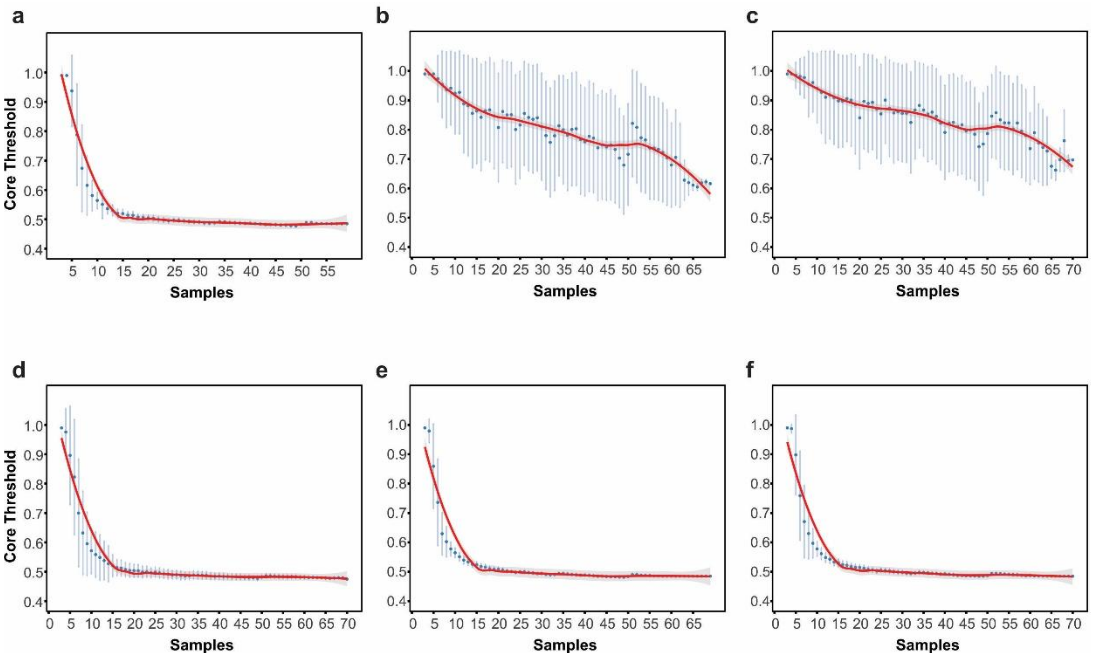


**Fig S2.** Relationship between the number of samples and the estimated core prevalence threshold across the compartments. a) Bacterial Leaves, b) Bacterial roots, c) Bacterial rhizosphere, d) ITS leaves, e) ITS roots and f) ITS rhizosphere. Blue points represent observed estimates, vertical bars indicate variability from subsampling, and the red line shows the fitted trend.

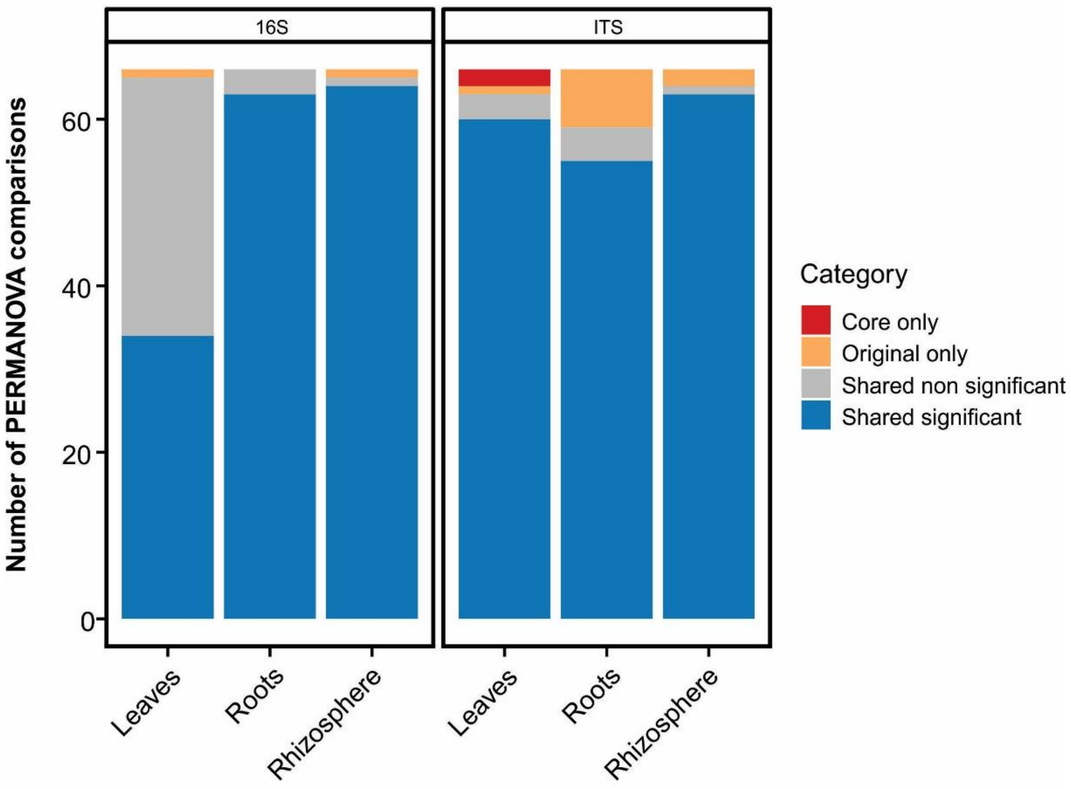


**Fig S3.** Summary of PERMANOVA significance consistency between original and core datasets across compartments and marker genes.

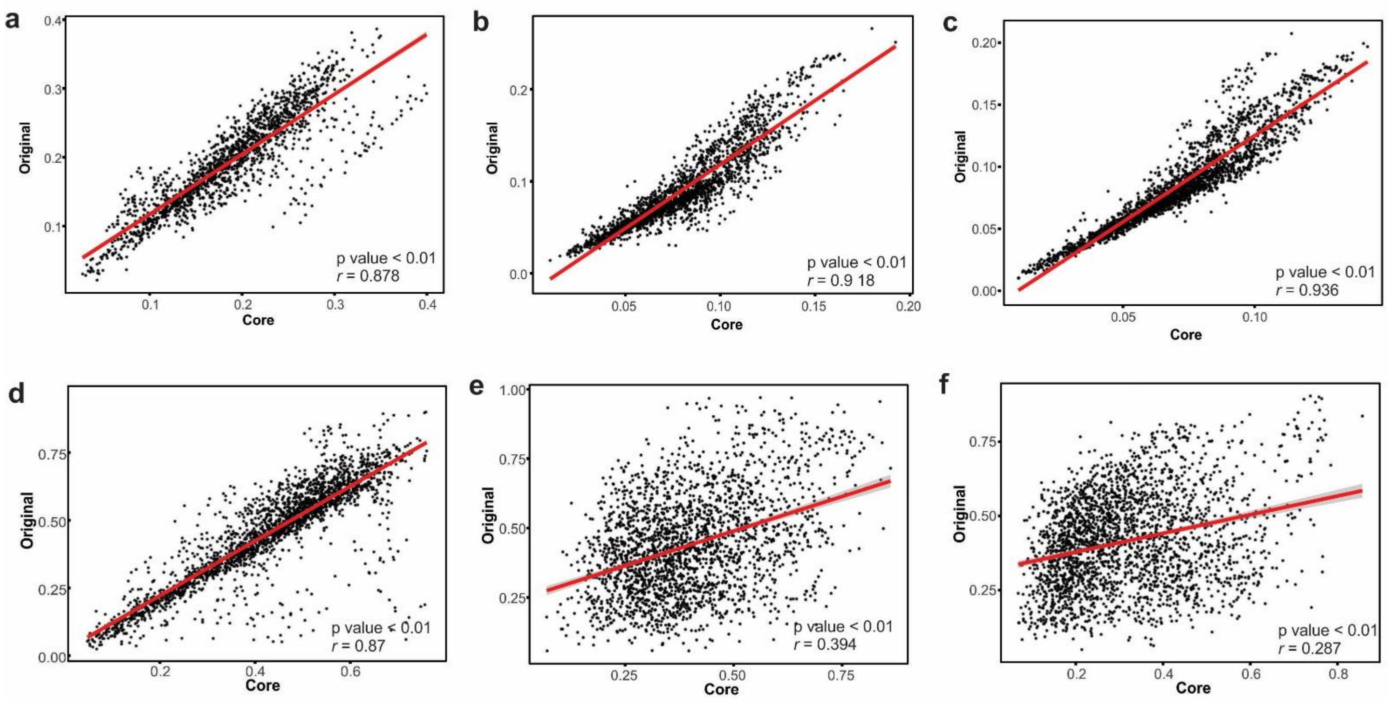


**Fig S4.** Mantel test correlation between the functional distance metric of the original and the core dataset. The top plots reparents the correlations between 16S matrix distances for a) leaves, b) roots and c) rhizosphere and the lower plots for the ITS matrix distances for d) leaves, e) roots and f) rhizosphere.

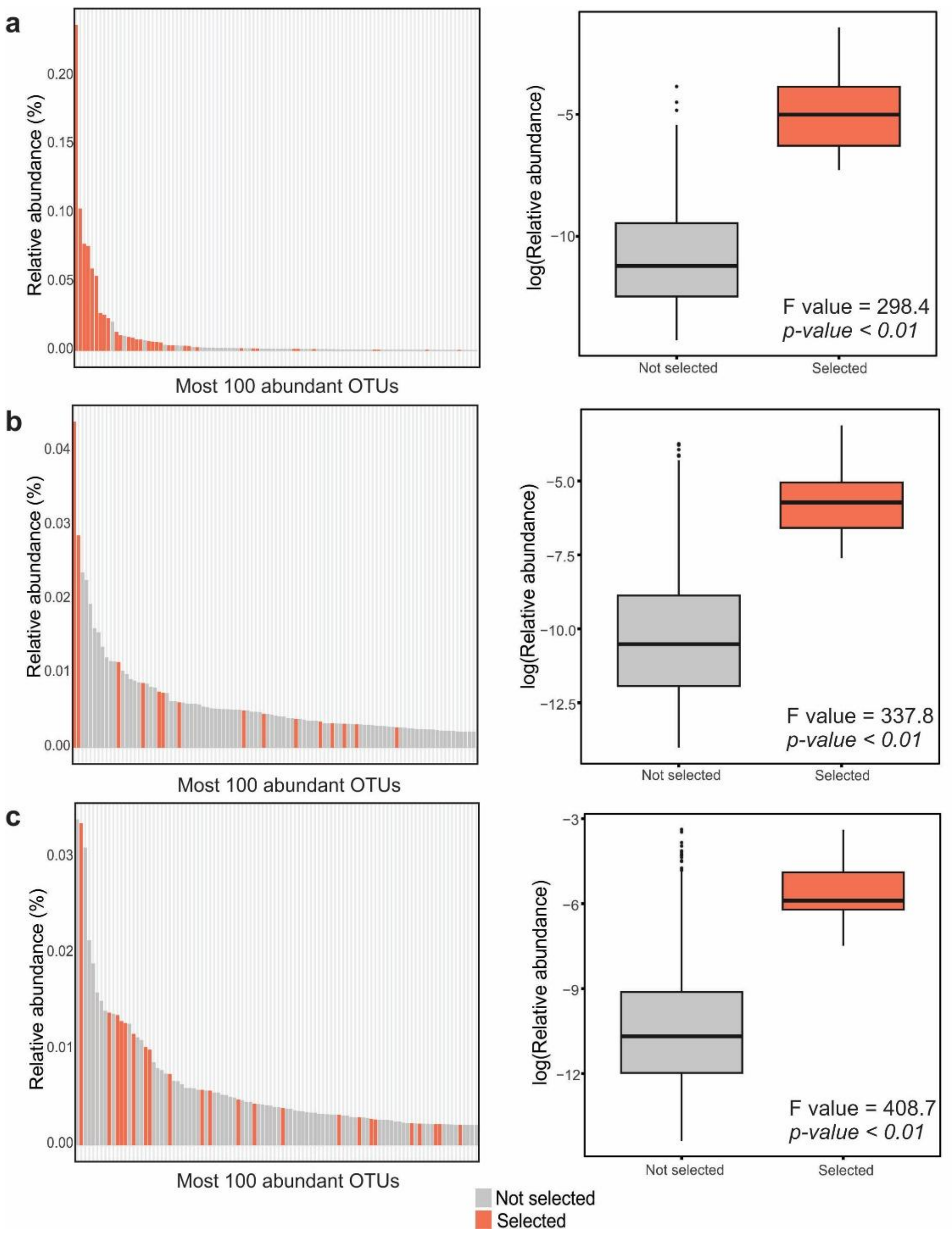


**Fig S5.** Relative abundance distribution of the 100 most abundant ITS OTUs across a) leaves, b) roots and c) rhizosphere. In the left panels, OTUs highlighted in red correspond to taxa selected by the core microbiome identification approach, whereas gray bars represent non-selected taxa. Right panels show boxplots comparing the log-transformed relative abundance of selected and non-selected OTUs.

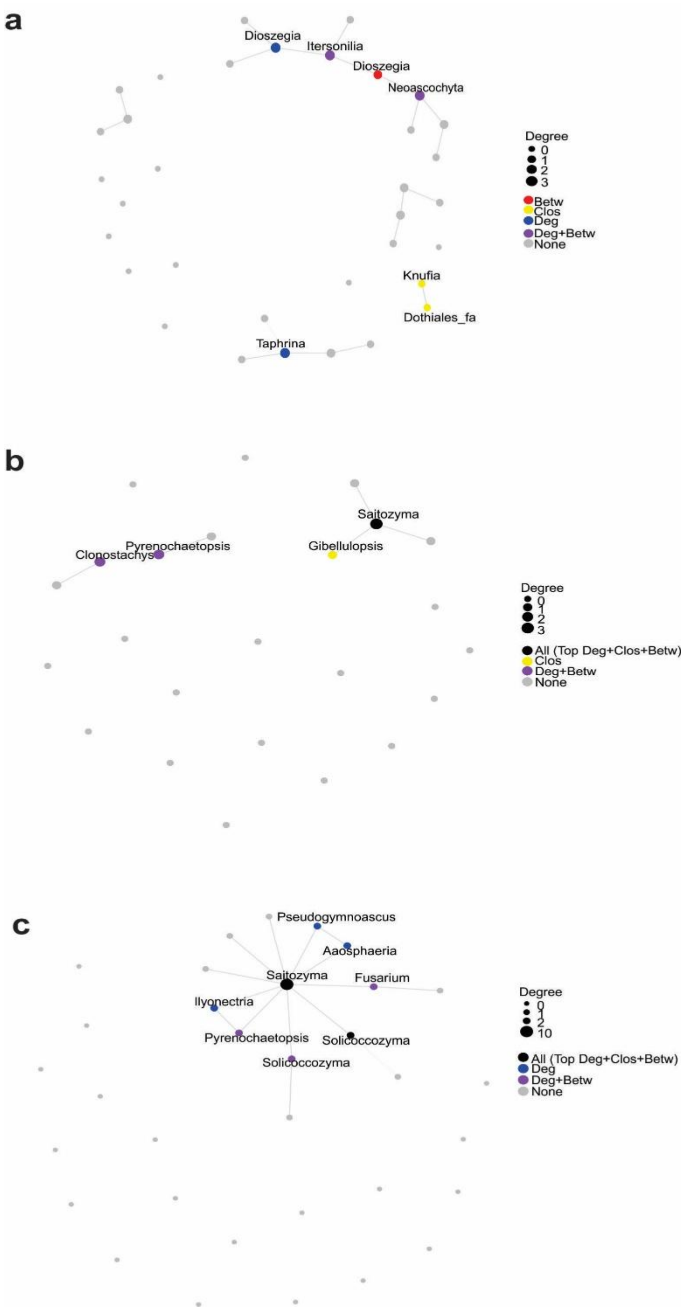


**Fig S6**. Network topology of the ITS microbial co-occurrence networks for (a) leaves, (b) roots, and (c) rhizosphere. Node size is proportional to degree (number of connections per node). Colored nodes highlight taxa identified as central according to different network centrality metrics: betweenness (Betw), closeness (Clos), degree (Deg). Gray nodes were not identified as central by any metric ("None"). Edges represent significant associations between taxa.
